# Targeting of NF-κB to Dendritic Spines is Required for Synaptic Signaling and Spine Development

**DOI:** 10.1101/283879

**Authors:** Erica C. Dresselhaus, Matthew C.H. Boersma, Mollie K. Meffert

## Abstract

Long-term forms of brain plasticity share a requirement for changes in gene expression induced by neuronal activity. Mechanisms that determine how the distinct and overlapping functions of multiple activity-responsive transcription factors, including nuclear factor kappa B (NF-κB), give rise to stimulus-appropriate neuronal responses remain unclear. We report that the p65/RelA subunit of NF-κB confers subcellular enrichment at neuronal dendritic spines and engineer a p65 mutant that lacks spine-enrichment (ΔSEp65) but retains inherent transcriptional activity equivalent to wild-type p65. Wild-type p65 or ΔSEp65 both rescue NF-κB-dependent gene expression in p65-deficient murine hippocampal neurons responding to diffuse (PMA/ionomycin) stimulation. In contrast, neurons lacking spine-enriched NF-κB are selectively impaired in NF-κB-dependent gene expression induced by elevated excitatory synaptic stimulation (bicuculline or glycine). We used the setting of excitatory synaptic activity during development that produces NF-κB-dependent growth of dendritic spines to test physiological function of spine-enriched NF-κB in an activity-dependent response. Expression of wild-type p65, but not ΔSEp65, is capable of rescuing spine density to normal levels in p65-deficient pyramidal neurons. Collectively, these data reveal that spatial localization in dendritic spines contributes unique capacities to the NF-κB transcription factor in synaptic activity-dependent responses.

**SIGNIFICANCE STATEMENT:** Extensive research has established a model in which the regulation of neuronal gene expression enables enduring forms of plasticity and learning. However, mechanisms imparting stimulus-specificity to gene regulation, insuring biologically appropriate responses, remain incompletely understood. NF-κB is a potent transcription factor with evolutionarily-conserved functions in learning and the growth of excitatory synaptic contacts. Neuronal NF-κB is localized in both synapse and somatic compartments, but whether the synaptic pool of NF-κB has discrete functions is unknown. This study reveals that NF-κB enriched in dendritic spines (the postsynaptic sites of excitatory contacts) is selectively required for NF-κB activation by synaptic stimulation and normal dendritic spine development. These results support spatial localization at synapses as a key variable mediating selective stimulus-response coupling.

## INTRODUCTION

Elucidating the mechanisms that allow cells to make stimulus-appropriate responses remains one of the most compelling questions in the field of gene expression. While cell-type specific transcription is heavily influenced by promoter accessibility, factors conferring stimulus-specificity to transcription have remained more elusive. Post-mitotic neurons in the adult brain are capable of differential activity-dependent gene regulation in response to seemingly subtle differences in stimuli patterns, magnitude, and duration. Mechanisms that precisely regulate gene target specificity dictate changes in the complement of neuronal proteins and, ultimately, such critical fates as whether neurons live or die, and whether synaptic connections are appropriately strengthened, weakened, or eliminated during processes such as information storage. Signaling cascades triggered downstream of neuronal activity by neurotransmitters, growth factors, and cytokines can individually and cooperatively induce multiple activity-responsive transcription factors, including CREB, Npas4, MEF2, MeCP2 (Nonaka et al., 2014), and nuclear factor-kappaB (NF-κB) (Shrum and Meffert, 2008). How these discrete transcription factors contribute unique or overlapping functions to disambiguate incoming signals and to orchestrate context appropriate changes in gene expression remains incompletely understood.

NF-κB transcription factors exhibit evolutionarily-conserved requirements in a wide variety of learning and memory paradigms (Freudenthal and Romano, 2000; Meffert et al., 2003; Freudenthal et al., 2005; Kaltschmidt et al., 2006; O’Riordan et al., 2006). NF-κB functions as a homo- or heterodimer of five possible mammalian subunits: p65 (also termed RelA), RelB, c-Rel, p50, and p52. The p65:p50 heterodimer is the most common NF-κB dimer in mammalian brain. Preformed NF-κB dimers are held latent in the cytoplasm by the NF-κB inhibitor, IkappaB (IκB) until stimulus-induced phosphorylation and degradation of IκB allows stable nuclear translocation of active NF-κB. NF-κB is present in neuronal somatic and dendritic cytoplasm and has been reported at synapses in both the *drosophila* neuromuscular junction (Heckscher et al., 2007) and in the mammalian CNS (Kaltschmidt et al., 1993) where synaptic NF-κB can be activated by excitatory stimulation (Meffert et al., 2003). Consistent with these results, components of the NF-κB activation pathway are also localized at synapses (Meffert et al., 2003). Previous work by our lab showed that NF-κB can be enriched in the postsynaptic compartment of dendritic spines from hippocampal neurons (Boersma et al., 2011), however, the potential for this localization to impart specific roles in activity-dependent responses was unknown.

In this study, we demonstrate that the p65 subunit of NF-κB is required for enrichment of the p65:p50 NF-κB dimer at dendritic spines and utilize a mutant p65 engineered to prevent spine-enrichment (p65ΔSE) to probe potential functions of NF-κB subcellular localization near neuronal synapses. Our results show that neurons lacking the spine-enriched pool of NF-κB are selectively deficient in NF-κB-dependent gene expression by stimuli delivered through excitatory synapses, while remaining competent to respond to stimulation that is not incoming through synapses. We assess a biological role for the spine-enriched pool of NF-κB using a model of hippocampal developmental synaptogenesis in which ongoing activation of NF-κB by elevated basal excitatory neurotransmission (Meffert et al., 2003; Mihalas et al., 2013) is required for appropriate dendritic spine formation (Boersma et al., 2011). In p65-deficient neurons, expression of wild-type p65, but not p65 lacking spine-enrichment (p65ΔSE), can rescue normal dendritic spine density in developing hippocampal neurons. Collectively, these findings reveal that subcellular localization is a salient feature in producing stimulus-specificity for activity-dependent gene expression and reveal a selective physiological function for the spine-enriched pool of NF-κB transcription factor in response to excitatory transmission and in dendritic spine development.

## MATERIALS and METHODS

### Animals and Primary Cultures

The care and use of mice met all guidelines of the local Institutional Animal Care and Use Committee (The Johns Hopkins University School of Medicine).

Hippocampal neurons were dissociated from post-natal day 0 (P0) wild-type (ICR), RelA^F/F^, or GFPRelA^F/F^ (De Lorenzi et al., 2009) male and female mouse pups and prepared as described in (Boersma et al., 2011). Cultures were transfected as described in (Boersma et al., 2011).

### Imaging and quantification

Confocal images of live murine hippocampal neurons were acquired on a Zeiss confocal microscope (LSM5 Pascal), and analyzed as described (Boersma et al., 2011). Briefly, fluorescence intensity in regions of interest (ROI) within dendritic spines and adjacent dendritic shafts from Z-stacks containing the entire neurons or process of interest were measured (ImageJ software) and analyzed using the following equations: Normalized spine fluorescence (NSF) = (EGFP_spine_/mCherry_spine_) / (EGFP_dendrite_/mCherry_dendrite_)

Percentage change in fluorescence = [NSF(p65tagged)-NSF(nontagged fluorescent protein)] / NSF(nontagged fluorescent protein)

Spine enrichment for endogenous GFPp65 was analyzed using expressed mCherry to mask the dendrite of interest (Imaris Bitplane software) and the spot function to place and quantify fluorescence for mCherry and GFP in spine-head ROIs.

### Neuronal Stimulation

DIV 20 - 21 hippocampal neuronal cultures from RelA^F/F^ mice (+ OHT) expressing either full length GFPp65 or GFPp65ΔSE were treated with synaptic or diffuse global stimuli at 37°C for 3.5 hours prior to harvest, as follows. For diffuse stimulation cultures were treated with Ionomycin (2 µM, EMD, 407951) and phorbol 12-myristate 13-acetate (PMA, 50 ng/ml, LC Labs, P-1680). For synaptic stimulation, neuronal cultures were either treated with bicuculline or with glycine in a low-dose version of chemical LTP. Bicuculline methobromide (30 µM, Tocris, 0109) was delivered for 30 seconds followed by 1x stop solution (100x stock: 1 mM kynurenic acid, 1 M MgCl2, 0.3 M NaOH, 500 mM NaHEPES, pH 7.3). For low-dose glycine stimulation, neuronal cultures were incubated with solution A (ACSF (130 mM NaCl, 3 mM KCl, 1.8 mM CaCl2, 2 mM MgCl2, 10 mM HEPES pH7.4, 10 mM D-Glucose), 1 μM Strychnine, 0.5 μM TTX (Tocris Bioscience)) for 10 minutes at 37°C, followed by 10 minutes with solution B (ACSF-0Mg + 150 μM Glycine, 1 μM Strychnine, 0.5 μM TTX) and then with solution A again for 3.5 hours at 37°C. Two distinct IKK (IκB kinase) inhibitors were used, where indicated. The selective IKK inhibitor TPCA-1 (2-[(Aminocarbonyl)amino]-5-(4-fluorophenyl)-3-thiophenecarboxamide, Tocris, 4 μM) or its vehicle, DMSO (dimethyl sulfoxide), or a TAT peptide coupled membrane-permeant nemo binding domain (NBD) peptide (ENZO, ALX-163-011) or control TAT peptide (ENZO, ALX-168-R050), both at 20 µM. IKK inhibitors were pre-incubated with neuronal cultures for 30 minutes prior to stimulation.

### Expression constructs and lentiviral preparation

Human p65, and p50 constructs were cloned by in-frame insertion using HindIII and BamHI into the Clontech C1 vector at the C terminus of enhanced green fluorescent protein (eGFP). p65 truncation mutants were created by PCR. Deletion mutants of p65 were created by two-step PCR.

PCR Primers:

p65Δ1-298: F-GTACGTAAGCTTCTGAGAAACGTAAAAGGACATATG,

R-GTACGTGGATCCTTAGGAGCTGATCTGACT

p65Δ305-406: F1-GTGCAAGCTTTAGACGAACTGTTCCCCCTCATCTTCCCG,

R1-CGTAGGATCCAAGGAGCTGATCTGACTCAGCAGGGCTGA,

F2-AAACGTAAAAGGACAGCCCCAGCCCCTGTCCCAGTC,

R2-GACAGGGGCTGGGGCTGTCCTTTTACGTTTCTCCTCAATCCGGTG

p65Δ335-442: F1-GTGCAAGCTTTAGACGAACTGTTCCCCCTCATCTTCCCG,

R1-CGTAGGATCCAAGGAGCTGATCTGACTCAGCAGGGCTGA,

F2-AAACGTAAAAGGACAGCCCCAGCCCCTGTCCCAGTC,

F2-ATTGCTGTGCCTTCCCAGCTGCAGTTTGATGATGAAGACCTGGGG,

R2-ATCAAACTGCAGCTGGGAAGGCACAGCAATGCGTCG

Lentivirus production from eBFP2-CreER^T2^ and eGFP2-CreER^T2^ were as described (Boersma et al., 2011; Mihalas et al., 2013). Hippocampal cultures from RelA^f/f^ mice were infected with CreER^T2^-expressing lentivirus at DIV 2 and treated with tamoxifen (OHT; 400nM, Sigma-Aldrich H6278) to elicit recombination for 4 days prior to experimentation.

### Reporter assays

Luciferase reporter assays were performed as described (Mihalas et al., 2013) by co-expressing firefly luciferase under the control of a promoter containing one or three NF-κB consensus binding sites in HEK293T and neurons, respectively, together with constitutively-expressed β-galactosidase (from pEF-Bos-LacZ), which was used to normalize for transfection efficiency and extract recovery.

### Immunocytochemistry and immunoblotting

Hippocampal neuronal cultures from GFPRelA^F/F^ mice were fixed at DIV 16 as described (Boersma et al., 2011) and subjected to immunohistochemistry with the following primary antibodies: chicken anti-GFP (AVES, GFP-1020,1:1000), rabbit anti-DsRed (Clontech, 632496, 1:500), and with secondary antibodies FITC anti-chicken (GeneTex, GTX77185, 1:500) and Alexa Fluor 568 anti-rabbit (Invitrogen, A11011,1:500) for 1 hour at room temp. in 10% BSA. Cells were mounted in 0.1M n-propyl gallate in 50% glycerol.

For immunoblotting, equivalent total protein levels (by BCA protein assay) from lysates were resolved by 12.5% SDS-PAGE gel electrophoresis followed by transfer to PVDF membrane and probed using: anti-p65 (Santa Cruz Biotechnology, sc372, 1:5000), anti-GFP (Neuromab, N86/8, 1:500 or Molecular Probes, A11122, 1:2000), anti-HSC70 (Santa Cruz Biotechnology, sc-7298, 1:10000).

### Electromobility shift assay

EMSAs were carried out using nuclear or cytoplasmic extracts prepared from isolated synaptosomes essentially as described (Meffert et al., 2003) and were reproduced 4 times. Extracts were incubated with a radiolabeled DNA oligonucleotide probe containing the κB sequence (wild-type: GGGGACTTTCC, mutant: GTTGACTTTCC).

### Statistical analyses

Graphs illustrate the arithmetic mean and error bars are SEM. For statistical analyses, unless otherwise noted two-tailed, unequal-variance *t* tests were used with α = 0.05. For ANOVA, one-way ANOVA with Bonferroni-Holm correction or Dunnetts correction was used. For unequal sample sizes, a Kruskal-Wallis test followed by posthoc Dunn’s multiple comparison tests was performed where indicated.

## RESULTS

### The p65 subunit of NF-κB is sufficient for localization to isolated synapses

We previously demonstrated that the p65 subunit of NF-κB is present primarily as a p65:p50 heterodimer in synapses isolated from murine hippocampi (synaptosomes), and that genetic deletion of p65 led to the absence of NF-κB from these synapses (Meffert et al., 2003). Multiple functional dimers of NF-κB exist in mammalian neurons, including p65:p50, p50:p50, and p65:p65 in the murine hippocampus. To investigate the subunit requirements for localization of NF-κB to neuronal synapses, we initially evaluated the effect of loss of the p50 subunit on the presence of NF-κB in isolated synapses. Biochemically isolated hippocampal synapses (synaptosomes) from mice wild-type for p50 or lacking the p50 subunit of NF-κB were subjected to electromobility shift assays (EMSA) with a radiolabeled DNA probe containing the κB sequence and antibody supershifts used to identify discrete NF-κB subunits. EMSA detects activated NF-κB dimers that are not IκB-bound and so are able to bind and shift the mobility of the DNA probe. Deoxycholate (DOC), to force dissociation of IκB, was included in some samples to reveal any occult (IκB bound) NF-κB, so that we would not fail to identify present but inactive forms of NF-κB (Meffert et al., 2003). We observed that isolated synapses from p50-deficient mice retained NF-κB consisting of p65:p65 homodimers as shown by the supershifted band in the presence of p65 antibody (Figure 1A; representative EMSA from 4 biological replicates). p65:p65 homodimers were present in increased abundance in the absence of the p50, in comparison to synapses isolated from wild-type neurons which contained predominantly p65:p50 heterodimers and small amounts of p65:p65 homodimers (Figure 1A). These results were consistent with p50 (when present) as the known preferred subunit binding-partner of p65, but demonstrated that, in the absence of p50, p65:p65 homodimers could localize to synapses. Since NF-κB is not detected in synapses isolated from p65-deficient neurons (Meffert et al., 2003), these results are collectively consistent with the p65 subunit being necessary as well as sufficient for significant localization of NF-κB to isolated hippocampal synapses.

**Figure 1:**
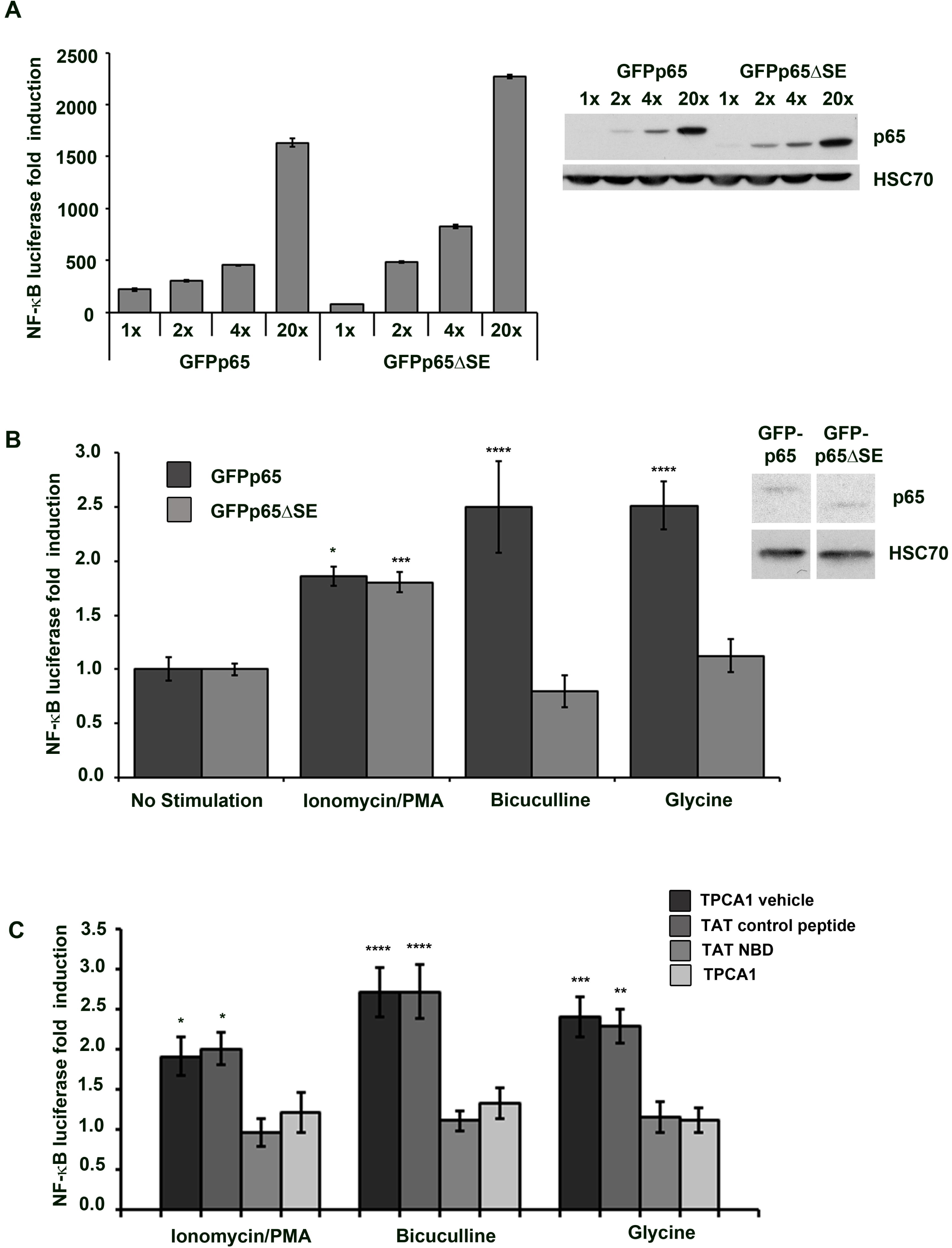
p65 is sufficient for NF-κB enrichment in dendritic spines of hippocampal pyramidal neurons. A) EMSA from synapses isolated from wild-type (p50^+/+^) or p50-deficient (p50^-/-^) murine hippocampi. NF-κB was detected using a radiolabelled oligonucleotide κB probe and specific subunits identified by antibody-supershifted bands. DOC was used to allow detection of occult (inactive) NF-κB. p65:p65 homodimers detected in p50^-/-^ isolated synapses from all (4) biological replicates of the experiment. B) Confocal images of live DIV17 hippocampal neurons co-expressing mCherry with either GFP, GFPp65 or GFPp50, as indicated. Graph shows % enrichment of GFPp65 or GFPp50, relative to GFP, as described in Materials and Methods. Scale bar = 5 µm. Error bars show SEM. \**p* = 3.8 x 10^-10^. C) Diagram representing spine measurements; dashed lines indicate diameter. D) Analysis of spine maturity from dendrites expressing GFPp65. Percent enrichment of GFPp65 in dendritic spines is calculated and binned according to spine size. Spine size was determined by a ratio of the spine head diameter to the spine neck diameter. Scale bar = 1 µm. Error bars show SEM. \**p*-values as compared to enrichment for spines <1 are as follows: for 1-3 *p* = 0.004, for 3-4 *p* = 0.002, for >4 *p* = 0.003. E) Confocal images of hippocampal pyramidal neurons co-expressing GFP and mCherry (left) or expressing mCherry fluorescent protein and GFPp65 from the endogenous RelA locus (right) and immunostained for GFP and mCherry. Graph shows percent enrichment of GFP and endogenously expressed GFPp65. Scale bar = 5 µm. Error bars show SEM. \**p* = 1.4×10^-5^.

### The p65 subunit of NF-κB, but not the p50 subunit, is enriched within dendritic spines

While biochemically isolated synapses can contain both pre-and post-synaptic elements, our previous work suggested that NF-κB was enriched in dendritic spines, the postsynaptic sites of excitatory synaptic contacts. Green fluorescent protein-tagged p65 (GFPp65) was enriched in spines of intact wild-type hippocampal neurons (Boersma et al., 2011) in which the p65:p50 dimer of NF-κB predominates (Meffert et al., 2003). To investigate whether p65 or p50 subunits were responsible for dendritic spine enrichment, we first excluded the possibility of p50 heterodimerization with p65, by assaying p50 spine-enrichment in neurons lacking p65. Hippocampal neuronal cultures from mice harboring a conditional loss of function allele for the RelA gene (encoding p65 protein, RelA^F/F^) were transduced with lentiviral inducible Cre recombinase (CreER^T2^) to generate p65-deficient cultures following administration of 4-hydroxy-tamoxifen (OHT). p65-deficient cultures co-expressing non-tagged mCherry fluorescent protein (for visualization and quantification) with either GFP-tagged p50 (GFPp50), GFPp65, or GFP alone, were subjected to confocal imaging and analysis for spine enrichment (calculation described in Materials and Methods). In the absence of endogenous p65, no spine enrichment of GFPp50 was observed (0.9 ± 1.2%), while GFPp65 retained its expected enrichment (39.7 ± 2.08%, *p* = 3.8 x 10^-10^) relative to GFP (Figure 1B). These results indicate that the p65 subunit of NF-κB is sufficient for hippocampal spine enrichment, and that p65 likely mediates spine enrichment of the predominant p65:p50 heterodimer.

### Characterization of spine parameters for p65 enrichment

Variation in the morphological parameters of dendritic spines, including size and head-to-neck ratio, have been shown to predict features such as spine stability and the presence of functional synapses (Nimchinsky et al., 2002; Holtmaat and Svoboda, 2009). In multiple brain regions, including the hippocampus, larger spine head diameter relative to neck diameter correlates with increased spine maturity and functional synapses, in comparison to stubby spines or spines lacking a head (Peters and Kaiserman-Abramof, 1970; Harris et al., 1992; Hering and Sheng, 2001; Tada and Sheng, 2006). We next investigated the possibility of a relationship between spine morphology and the extent of p65 enrichment in dendritic spines. Confocal Z-stacks of dendritic spines from hippocampal neurons co-expressing GFPp65 and mCherry were analyzed to calculate the ratio of dendritic spine head diameter to neck diameter (Figure 1C), as well as spine enrichment of GFPp65. Spines were considered any protrusion between 0.2 µm to 3.0 µm with or without a head. For comparison and based on published characterizations, spines were classified into 4 categories: spines in which the head to neck ratio was ≤ 1, between 1 - 3, between 3 - 4, or > 4. Spines in which the head to neck diameter ratio was ≤ 1 (i.e. lacking a distinct head) did not exhibit significant enrichment of GFPp65 at the spine terminus (GFPp65 spine enrichment of 7.99 ± 6.04%, *p* = 0.056 compared to GFP alone) (Figure 1D). In contrast, all three spine categories with head to neck diameter ratio ≥ 1 had significantly more GFPp65 spine enrichment than spines with head to neck ratio ≤ 1 (*p* ≤ 0.003, ANOVA *p* = 0.005, Dunn test for nonparametric data *p* ≤ 0.01). A trend was observed for greater GFPp65 enrichment with increasing spine head to neck ratio: head to neck ratio 1-3 (average enrichment 33.57 ± 2.47%), head to neck ratio 3 - 4 (average enrichment 41.41 ± 6.31%), head to neck ratio > 4 (average enrichment 46.81 ± 9.34%) (Figure 1D), but this did not reach significance (ANOVA, *p* = 0.159). These data indicated that GFPp65 was more enriched in head-containing spines with a greater likelihood of mature synaptic connections and suggested the possibility of a role for NF-κB at functional excitatory synapses.

### p65 expressed from the endogenous locus is enriched in dendritic spines

While expression of exogenous constructs was routinely titrated to the lowest possible levels, we sought to further test the possibility of inappropriate p65 localization due to overexpression artifact prior to further investigation. Anti-p65 antibodies competent for immunoblotting have been identified, however, antibodies capable of reliable brain immunostaining without staining in p65-deficient brain tissue were not identified despite candidate screening. For this reason, we proceeded to assess whether expression from the endogenous locus of p65 would similarly exhibit p65 enrichment in dendritic spines using a homozygous transgenic knock-in mouse line expressing the fusion protein of GFP and p65 (GFPp65) from the genomic RelA locus with GFP inserted immediately following the start codon (GFPRelA^f/f^) (De Lorenzi et al. 2009). Notably, characterization of this line demonstrates GFPp65 expression to be equal to endogenous p65 expression in wild-type mice, and to fully rescue endogenous p65 function (p65 loss is embryonic lethal).

Hippocampal cultures from GFPRelA^f/f^ mice were transfected with mCherry to permit morphological isolation of individual neurons and enrichment quantification, and subjected to immunostaining for GFP and mCherry (see Materials and Methods), followed by confocal microscopy and 3-dimensional analysis of Z-stack projections (Imaris). p65 expressed from the endogenous locus was enriched in hippocampal dendritic spines by 33.5 ± 6.9% (*p* = 1.36×10^-5^) relative to non-tagged GFP (Figure 1E); this enrichment was not significantly different from the dendritic spine enrichment of transiently-transfected GFPp65 as shown in Figure 1B (*p* = 0.27). These results supported the use of GFPp65 constructs expressed at low levels in subsequent experiments to investigate the determinants of p65 enrichment.

### A p65 deletion mutant lacking dendritic spine-enrichment

We next engineered a series of deletion and truncation constructs (Figure 2A) aimed at identifying a mutant of the p65 subunit that would lack dendritic spine-enrichment and allow us to examine potential physiological roles of a synaptic pool of NF-κB. All NF-κB subunits share a highly homologous amino-terminal domain, known as the Rel Homology Domain (RHD), which contains sequences responsible for DNA binding, dimerization, IκB binding, and nuclear localization. p65 also contains a carboxy-terminal trans-activation domain (TAD), which is a shared feature of several NF-κB subunits (Napetschnig and Wu 2013). Amino-terminally GFP-tagged p65 truncations lacking either the carboxy-terminal TAD (p65ΔTAD) or the amino-terminal RHD (p65Δ1-298) were co-expressed with non-tagged mCherry fluorescent protein in hippocampal pyramidal neurons, subjected to confocal imaging, and found to retain dendritic spine enrichment that did not significantly differ from wild-type GFPp65 (40.8 ± 4.9%, *p* = 0.813 and 40.29 ± 3.95% *p* = 0.980 respectively,) (Figure 2B and 2C). In contrast, a p65 truncation consisting solely of the amino-terminal RHD including the NLS (GFPp65Δ305-551), lacked enrichment in dendritic spines and was not significantly different in enrichment compared to GFP (−4.2 ± 2.4%, *p* = 0.142). Collectively these data, prompted us to focus on the region between the NLS and the TAD of p65, a relatively uncharacterized region with little sequence conservation between NF-κB subunits, which we suspected might be involved in dendritic spine enrichment.

**Figure 2:**
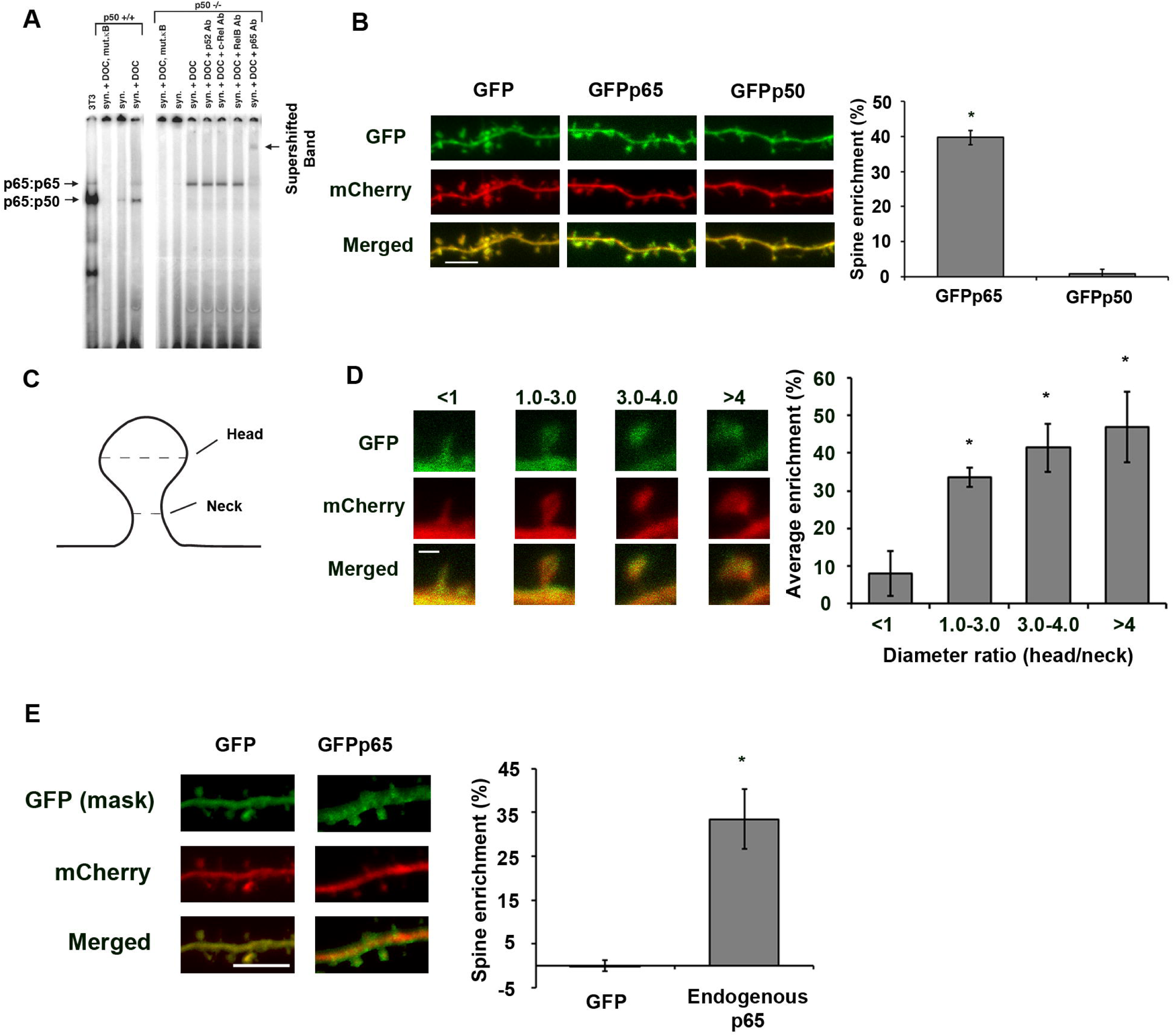
Identification of non-spine enriched mutants of p65. A) Diagram of p65 and p65 deletion and truncation constructs. B) Graph showing % enrichment of GFP-tagged p65 and GFP-tagged p65 deletion and truncation constructs. Enrichment is determined as described in Materials and Methods. Error bars show SEM. \**p*-values as compared to GFP condition are as follows: for GFPp65 *p* = 2.5×10^-30^, for GFPp65Δ1-298 *p* = 4.7×10^-15^, for GFPp65Δ335-442 *p* = 9.5×10^-8^, and # data from GFPp65ΔTAD condition was previously presented (Boersma et al., 2011) in a different format. C) Representative confocal images from live murine hippocampal neurons co-expressing mCherry with GFP-tagged p65 deletion or truncation constructs or wild-type p65. Scale bar = 5 µm. D) p65 constructs were expressed in 293T cells at equivalent levels and migrate at their expected molecular weights by immunoblot of lysates. Expected molecular weights (kDa) as follows: GFPp65: 87, GFPp65ΔTAD: 76, GFPp65Δ1-298: 53, GFPp65Δ1002-1326: 76, GFPp65Δ916-1218: 76, GFPp65Δ305-551: 62, GFP: 27.E) Expression of p65 constructs does not significantly alter measures of neuronal health. Percent of neurons scoring positive in confocal imaging for cytoplasmic blebbing/beading or for pyknotic nuclear morphology using Hoechst staining from hippocampal neurons co-expressing mCherry with either GFP, GFPp65, or GFPp65ΔSE. Staurosporine treatment of GFP-expressing neurons serves as a positive control; experimenter was blinded to condition during all imaging and analysis. Cells were scored as positive or negative (see Materials and Methods) and percent positive plotted per condition (precluding SEM calculation).

Two constructs with overlapping deletions in p65 between the NLS and TAD (p65Δ305-406 and p65Δ335-442) were engineered to refine the region required for enrichment. Equivalent expression of these GFP-tagged constructs in hippocampal pyramidal neurons co-expressing mCherry revealed that p65Δ335-442 retained significant enrichment in comparison to GFP alone, while enrichment remained lower than for full-length p65 (19.1 ± 3.5%, *p* = 4.67×10^-5^ and *p* = 0.002) (Figure 2B,C). In contrast, p65Δ305-406 lacked significant spine enrichment (5.5 ± 2.2%, *p* = 0.102 compared to GFP alone), and we termed this mutant, GFPp65ΔSE. All truncation and deletion p65 constructs were expressed at similar levels, as assessed by visualization of the GFP-tag in confocal imaging (Figure 2C), and yielded protein products of the expected molecular weights by immunoblotting of lysates from transfected 293T cells (Figure 2D).

Due to the known participation of NF-κB in apoptotic pathways, we evaluated GFPp65ΔSE expression for potential effects on neuronal cell health prior to proceeding to other assays of physiological function. Dysregulation of NF-κB signaling in some contexts can produce apoptosis which has been visualized as fragmented or pyknotic nuclei or as cytoplasmic blebbing or beading (Baichwal and Baeuerle, 1997). We assessed cytoplasmic integrity and nuclear morphology by confocal imaging of live hippocampal neurons expressing either GFP, GFPp65 or GFPp65ΔSE for 24 hours, together with mCherry fluorescent protein to fill the cytoplasm and Hoechst staining to visualize nuclei. The experimenter was blinded to condition during all imaging and analysis. Quantification showed no differences in cytoplasmic blebbing (GFP: 19.2%, n = 9 cells; GFPp65: 0%, n = 8; GFPp65ΔSE: 12.5%, n = 8) or pyknotic nuclei (GFP: 11.1%, n = 9 cells; GFPp65: 12.5%, n = 8; and GFPp65ΔSE: 12.5%, n = 8) between neurons expressing GFP, GFPp65, or GFPp65ΔSE (Figure 2E). In contrast, cytoplasmic blebbing and pyknotic, condensed and fragmented nuclei were readily detected in 63.6% and 72.7%, respectively, of control neurons (n = 11) expressing GFP and treated with the kinase inhibitor, staurosporine (50 nM, 12hr) to induce apoptosis (Figure 2E). These results indicated that GFPp65ΔSE expression did not adversely affect neuronal health as monitored by nuclear morphology and cytoplasmic blebbing/beading. We proceeded to assess the transcriptional function of GFPp65ΔSE, in comparison to wild-type p65.

### Non-spine enriched p65 mutant retains basal transcriptional activity

We first examined whether the inherent transcriptional properties of wild-type p65 were altered in GFPp65ΔSE. While the region of p65 between the RHD and TAD, containing the deletion in GFPp65ΔSE, has not been previously shown to contain domains implicated in transcriptional regulation, we directly assessed the inherent capacity of GFPp65ΔSE to induce NF-κB dependent transcription in comparison to wild-type p65 (GFPp65) by dose titration in an NF-κB reporter assay. This reporter assay effectively assesses multiple aspects of the NF-κB transcription factor, including appropriate protein folding, stability, dimerization, DNA-binding, and transactivation capacity. The relative inherent activity of an NF-κB construct is evaluated by this reporter assay in heterologous cells, without the need for cellular stimulation to mediate IκB inhibitor degradation, through moderate overexpression of NF-κB subunits which intentionally outstrips endogenous IκB to allow assessment of transcription activation. HEK293T cells were co-transfected with NF-κB luciferase reporter and a constitutively expressed β-galactosidase for normalization, in combination with either full length GFPp65 or GFPp65ΔSE at increasing doses which were titrated for equivalent levels of protein expression by immunoblot (Figure 3A, right). Full length GFPp65 and GFPp65ΔSE could each similarly activate NF-κB-dependent transcription from the luciferase reporter across a range of expression levels in the dose titration (Figure 3A, left). We conclude that deletion of amino acids 305-406 in p65ΔSE lacks measurable impact on the inherent capability of p65 to induce NF-κB-dependent transcription.

**Figure 3:**
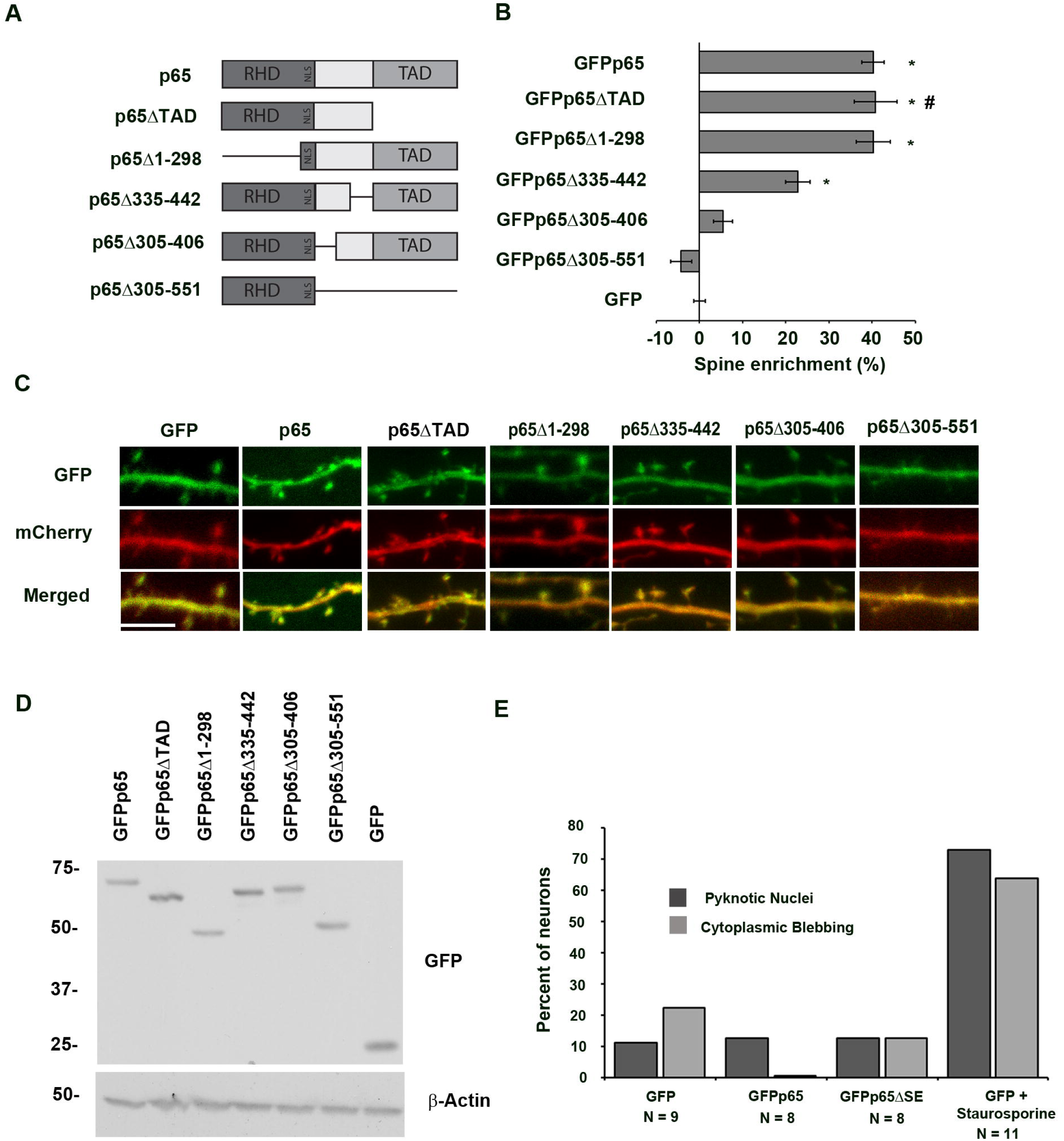
A p65 mutant lacking spine-enrichment is selectively deficient in response to synaptic stimulation. A) Inherent transactivation potential of wild-type p65 and p65 lacking spine-enrichment (p65ΔSE) are not distinguishable. Reporter assay for dose-titrated expression of GFPp65 or GFPp65ΔSE in HEK293T cells co-expressing NF-κB luciferase reporter and constitutive β-galactosidase used for normalization (Left). Fold induction calculated as described in Materials and Methods. Representative immunoblot of dose titration (Right). B) NF-κB luciferase assay of p65-deficient neurons expressing full length GFPp65 or GFPp65ΔSE and either mock-stimulated or stimulated with bicuculline (30 μM for 30 seconds), with ionomycin/PMA (3.5 hr), or with glycine (100 µM for 10 min). \**p* = 0.018 compared to GFPp65 no stimulation. \*\**p* = 0.0002 compared to GFPp65ΔSE no stimulation. \*\*\*\**p* = 0.0001 compared to GFPp65 no stimulation. For neurons expressing GFPp65ΔSE, NF-κB reporter activity in response to bicuculline or glycine does not differ significantly from the no stimulation condition. Representative immunoblot (inset) illustrating similar levels of expressed protein for GFPp65 or GFPp65ΔSE in these experiments. C) NF-κB luciferase assay of wild-type neurons either mock-stimulated or stimulated with ionomycin/PMA (2 µM and 50 ng/ml), bicuculline (30 μM for 30 seconds), or with glycine (150 µM for 10 min) and treated with either TPCA1 (4 µM) or TAT-NBD (20 µM) or their respective vehicle or TAT controls. \**p* = 0.04, \*\*\**p* = 0.0007, \*\*\*\**p* = 0.0001, all compared to no stimulation which was set to 1. Error bars indicate SEM.

### Non-spine enriched p65 mutant is not activated by a synaptic stimulus

We next conducted experiments in hippocampal pyramidal neurons aimed at addressing whether loss of spine enrichment might selectively alter NF-κB-dependent transcription in response to stimuli originating at the excitatory glutamatergic synapses which occur predominantly on dendritic spines, in comparison to stimuli with a diffuse origin. Following robust excitatory synaptic activation, suprathreshold ion fluxes or second messenger signals can traverse from dendritic spines to dendritic and somatic compartments (Nimchinsky et al., 2002). However, signals (including calcium elevations) initiated by lower and physiological levels of synaptic activity, can be either restricted to activated dendritic spines or near-neighbor spines (Nimchinsky et al., 2002; Noguchi et al., 2005), or diminished prior to reaching the neuronal soma (Andersen et al., 1980; Yuste et al., 2000). This prompted us to ask whether one role of a spine-localized pool of NF-κB might be to preferentially regulate gene expression in response to incoming stimuli in which signals are spatially restricted or of greater amplitude near excitatory synapses on dendritic spines. To test this hypothesis we examined whether modest excitatory synaptic stimulation could activate NF-κB lacking dendritic spine enrichment (GFPp65ΔSE) as effectively as wild-type NF-κB (GFPp65) which is spine-enriched. The participation of endogenous p65 was excluded by conducting NF-κB reporter assays in p65-deficient hippocampal cultures prepared from RelA^f/f^ mice. Mature neuronal cultures (DIV20-21, when basal excitatory activity is low) were co-transfected with NF-κB reporter and constitutively-expressed β–galactosidase, and low levels of either full-length GFPp65 or GFPp65ΔSE. GFPp65ΔSE and GFPp65 were expressed at similar levels as assessed by immunoblot (Figure 3B, inset) and fluorescence confocal imaging. Neurons were then treated with either stimuli which enhance endogenous excitatory synaptic transmission or with a diffuse stimulus not originating at synapses. Synaptic stimulation was delivered using low-dose bicuculline (30 μM for 30 seconds), a GABA_A_ receptor inhibitor that enhances endogenous glutamatergic excitatory transmission, or low-dose glycine (a chemical LTP protocol stimulating NMDA receptors only at synapses receiving spontaneously released glutamate)(Fortin et al., 2010; Araki et al., 2015). Calcium ionophore (ionomycin, 2 μM) and phorbol ester (PMA, 50 ng/ml) were used to deliver diffuse generalized stimulation. Both synaptic and diffuse stimuli have been previously characterized as activators of endogenous NF-κB (Meffert et al., 2003). In the absence of stimulation, neurons expressing low level GFPp65 or GFPp65ΔSE exhibited similar levels of normalized NF-κB reporter activity, and stimulations were graphed relative to this level (set as 1.0, Figure 3B). Ionomycin/PMA stimulation equivalently increased NF-κB reporter activity in neurons expressing either GFPp65 or GFPp65ΔSE (1.86 ± 0.09 and 1.80 ± 0.09 fold, *p* = 0.018 and *p* = 0.0002, respectively, compared to no stimulation). In contrast, synaptic activation with low dose bicuculline increased NF-κB reporter expression by 2.58 ± 0.39 fold in neurons expressing GFPp65 while neurons expressing GFPp65ΔSE showed no significant response (*p* = 0.0001 and *p* = 0.389, respectively, compared to no stimulation (Figure 3B). Synaptic stimulation with glycine increased NF-κB reporter expression by 2.52 ± 0.21 fold in neurons expressing GFPp65 while neurons expressing GFPp65ΔSE showed no significant response (*p* = 0.0001 and *p* = 0.692, respectively, compared to no stimulation (Figure 3B). Collectively, this data indicates that GFPp65ΔSE, which lacks enrichment at dendritic spines, can support NF-κB-dependent gene expression in response to a diffuse stimulus, but is ineffective in supporting NF-κB-dependent gene expression in response to two stimuli originating in the synaptic compartment. These findings are consistent with a role for the spine-enriched pool of NF-κB in mediating NF-κB-dependent gene expression in response to excitatory synaptic stimuli.

### Response to synaptic stimuli requires the canonical NF-κB pathway

Previous work has shown that signaling components required for NF-κB activation, including degradation of the inhibitor of NF-κB (IκB), are present at synapses (Meffert et al., 2003). The failure of synaptic stimulation to induce NF-κB-dependent transcription in neurons lacking spine-enriched NF-κB could be due to a necessity for localization of the NF-κB/IκB complex in proximity to stimuli incoming at synapses. To test whether synaptic and diffuse stimuli utilize distinct or shared NF-κB induction pathways, we assessed the effect of inhibition of the canonical activation pathway through IKK, using both small molecule (Skaug et al., 2011; Liu et al., 2012) and peptide IKK inhibitors(May et al., 2000; Solt et al., 2007; Solt et al., 2009). The selective small molecule IKK inhibitor, TPCA-1, or a cell-permeant nemo (IKKγ) binding domain IKK inhibitor peptide (TAT-NBD) each prevented increased NF-κB activation by diffuse or by synaptic stimuli (Figure 3C; Ionomycin/PMA, Bicuculline, or Glycine) in hippocampal neurons, compared to control stimulations in vehicle alone (Ionomycin/PMA: 1.91 ± 0.24, p = 0.043, Bicuculline: 2.72 ± 0.31, p = 0.0001, glycine: 2.41 ± 0.25, p = 0.0007) or control TAT peptide (Ionomycin/PMA: 2.01 ± 0.20, p = 0.014, Bicuculline: 2.72 ± 0.33, p = 0.0001, glycine: 2.30 ± 0.21, p = 0.002).). These data demonstrate that NF-κB activation by diffuse and synaptic stimuli both proceed through the canonical IKK-dependent pathway, consistent with a difference in localization of the NF-κB complex near synapses underlying the loss of responsiveness in neurons which were lacking spine-enriched NF-κB.

### Spine-enriched NF-κB in dendritic spine development

During developmental periods of rapid spine and synapse formation, high basal levels of excitatory glutamatergic synaptic transmission produce elevated NF-κB transcriptional activity which is required for the *in vitro* and *in vivo* production of normal hippocampal pyramidal dendritic spine density, spine maturity, and corresponding synaptic currents (Boersma et al., 2011; Schmeisser et al., 2012). We harnessed this assay to further probe function of the spine-enriched pool of NF-κB by testing its requirement in a biologically relevant readout, dendritic spine formation, downstream of synaptic stimulation. Hippocampal cultures deficient in endogenous p65 (from RelA^f/f^ mice) were transfected with either GFP, wild-type p65 (GFPp65), or the p65 mutant lacking spine-enrichment (GFPp65ΔSE) and pyramidal neurons assayed for dendritic spine density and spine head volume by live confocal imaging during the period of rapid spine formation (DIV 14 - 16) (Figure 4). Basal spine density in wild-type neurons expressing GFP was 1.89 ± 0.23 spines / 10 µm, with an average spine head volume of 0.217 ± 0.028 µm^3^. Loss of p65 (GFP + OHT) decreased spine density by 46.7 % (to 1.01 ± 0.08 spines / 10 µm) and average spine head volume by 40.4 % (to 0.129 ± 0.021 µm^3^). Expression of GFPp65 in p65-deficient neurons rescued both spine density (2.35 ± 0.27 spines / 10 µm) and spine head volume (0.258 ± 0.021 µm^3^) to levels indistinguishable from wild-type neurons. In contrast, dendritic spine density and volumes in p65-deficient neurons expressing GFPp65ΔSE did not significantly differ from neurons lacking p65 (GFP + OHT). Expression of GFPp65ΔSE in p65-deficient neurons also failed to rescue spine density (1.53 ± 0.20 spines / 10 µm, *p* = 0.03) to wild-type levels, and showed a corresponding trend toward failed rescue for spine head volumes (0.154 ± 0.020 µm^3^, *p* = 0.073). Endogenous p65 is required for the development of normal spine size and density. The inability of GFPp65ΔSE to mimic the effects of wild-type p65 on spine density and head volume indicates a critical biological role for spatial localization of p65 to dendritic spines in supporting spine growth, which relies upon NF-κB-dependent transcription (Boersma et al., 2011).

**Figure 4:**
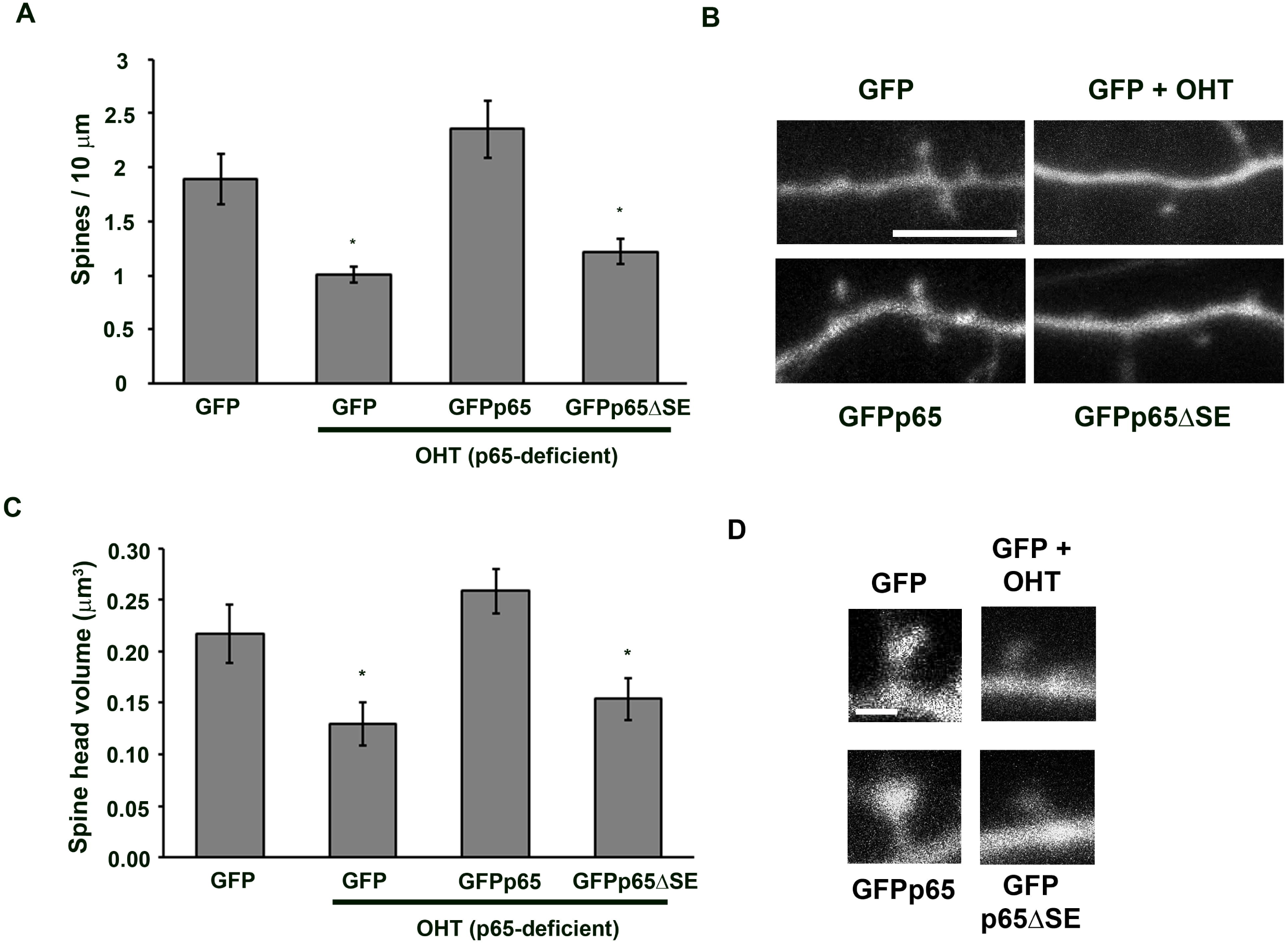
p65 lacking spine-enrichment fails to rescue neuronal dendritic spine density and size during spinogenesis. Loss of endogenous p65 (OHT) in pyramidal neurons (DIV 15-16) significantly decreases dendritic spine density (GFP + OHT, \**p* = .0047, panel A) and spine head volume (GFP + OHT, \**p* = 0.0151, panel C), compared to GFP. A) Expression of wild-type p65 (GFPp65, *p* = 0.212), but not p65 lacking spine enrichment (GFPp65ΔSE, *\*p* = 0.024), rescues dendritic spine density back to baseline (GFP) levels in hippocampal pyramidal neurons conditionally deficient in p65 (OHT) (ANOVA, *p* = 0.0009). B) Representative confocal projections (with co-expressed mCherry) of dendrites from live murine hippocampal neurons used to calculate spine densities (A). Scale bar = 5 µm. C) Expression of wild-type p65 (GFPp65, *p* = 0.262) rescues spine head volume back to baseline (GFP) levels in p65-deficient hippocampal pyramidal neurons. Expression of GFPp65ΔSE shows a trend towards failed rescue of spine head volume (*p* = 0.073, compared to GFP) (ANOVA, *p* = 0.0056) and spine head volumes which are significantly different from GFPp65-expressing neurons (#, *p* = 0.0006, compared to GFPp65 + OHT). D) Representative confocal projections (with co-expressed mCherry) from live murine hippocampal neurons used to calculate spine head volumes (C). Scale bar = 1.0 µm. Error bars indicate SEM.

## DISCUSSION

Healthy neural plasticity in the adult brain is enabled by the selective regulation of gene expression following neuronal activity. Features that enable transcription factors to differentiate neuronal stimuli, and to produce appropriate gene regulatory responses are incompletely understood. In this work, we show that the enrichment of a transcription factor in a discrete subcellular location, the dendritic spine head, selectively facilitates responses to excitatory activity incoming through synapses, while not being required for the response to more diffuse neuronal stimulation that does not originate at synapses.

NF-κB is a pleiotropic transcription factor that functions broadly in the control of genes promoting cellular and synapse growth, and for which evolutionarily conserved requirements have been established in multiple assays of plasticity, learning, and memory from crabs to fruit flies to mammals (Shrum and Meffert, 2008; Salles et al., 2014). The characterization of NF-κB enrichment in dendritic spines presents an interesting distinguishing feature, as relatively few transcription factors are known to localize to discrete extranuclear regions. Other notable exceptions include CREB (which has been reported in axons), and Stat3 and the transcription activator ELK-1 which have been reported in neuronal dendrites (Suzuki et al., 1998). We identify a 101 amino acid region of the p65 NF-κB subunit that is required for NF-κB enrichment in dendritic spine heads. Overlapping mutants (Figure 2) suggest that the region of importance for dendritic spine localization may be further narrowed to a 30 amino acid sequence. No protein-protein or protein-RNA interaction motifs have been previously characterized in this region of p65, which bears little conservation among other NF-κB subunits. A motif scan (Scansite 4.0, MIT) does identify a Src homology 3 (SH3) domain poly-proline binding motif in p65 which is conserved across mammals (beginning at human P322, PRPPP), and is lacking in GFPp65ΔSE but present in all spine-enriched constructs of p65 (Figure 2A,B). Binding of SH3 domains has been implicated in mediating protein-protein interactions and clustering at synapses; for example, MAGUK scaffolding proteins contain SH3 domains and are concentrated in the postsynaptic densities of neuronal synapses (Sheng, 1996; McGee and Bredt, 1999). In previous work (Boersma et al., 2011), we demonstrated that a MAGUK, PSD-95, is an NF-κB transcriptional target that is critical for NF-κB-mediated increases in dendritic spine density. Additional analysis of the amino acid region implicated in spine-enrichment of p65 by charge density showed that it contained the entirety of a long stretch of uncharged amino acids (82 amino acids). When subjected to analysis for protein disorder prediction (DISOPRED3, PSIPRED protein sequence analysis UCL; (Jones and Cozzetto, 2015)) the p65 profile showed that the section implicated in spine enrichment (amino acids, 305 – 406), was predicted with high confidence to contain an intrinsically disordered region (IDR, lacking defined and ordered 3D structure). This is particularly intriguing given the recently heightened interest in potential biological roles of protein IDRs (Lin et al., 2015; Shin and Brangwynne, 2017; Wei et al., 2017).

Biochemical and ionic fluxes between the cytoplasm of spines and neighboring dendritic shafts can be limited in mature spines, many of which have relatively thin necks connected to the parent dendrite (Bourne and Harris, 2008; Adrian et al., 2014). This type of compartmentalization may confer particular value to spine-enrichment of NF-κB in sensing synaptic excitation and mediating activity-responsive gene expression in neurons. Previous work has shown that neuronal NF-κB can be activated by sub-membranous calcium elevation signaling through calcium-calmodulin protein kinase II (CaMKIIα, a kinase highly abundant in the post-synaptic density), in hippocampal neurons as well as in isolated synapses (Meffert et al., 2003). In this report, we demonstrate a role for local enrichment in dendritic spines in shaping the stimulus specificity of NF-κB. The p65 mutant engineered to lack spine head enrichment, p65ΔSE, retains inherent NF-κB transcriptional activity and transcriptional response to diffuse stimulation, but exhibits deficiency in mediating NF-κB-dependent transcription to modest stimulation incoming through excitatory synapses and fails to support normal head volume and density of dendritic spines. The enrichment of NF-κB in dendritic spine heads raises the possibility of a potential contribution of local protein-protein interactions not requiring transcription to these phenotypes. However, using transcriptionally-inactive mutants of p65 (including p65ΔTAD), we previously found that activity-dependent enhancement of dendritic spine size and density required cell-autonomous NF-κB-dependent transcription (Boersma et al., 2011).

NF-κB activation by either diffuse or synaptic stimulation required the canonical IKK-mediated pathway, previously shown to participate in synaptic plasticity (Russo et al., 2009). This shared feature further supports the importance of subcellular localization of the initiating stimulus in defining the requirement for spine-enriched NF-κB. It is possible that enrichment at the sites of incoming excitatory stimuli might also regulate additional features of gene expression, by impacting the kinetics, magnitude, or duration of NF-κB induction following stimulation. Since the exact sequence of the NF-κB DNA-response elements in a promoter/enhancer region is reported to code a preference for binding to (and activation by) particular NF-κB dimers under some conditions (Wang et al., 2012), selective activation of p65-containing NF-κB dimers that are spine-enriched might have the potential to impact the specificity of NF-κB-dependent gene expression. Collectively, this work highlights spine head enrichment as a feature contributing to activity-responsive gene expression by NF-κB, a transcription factor implicated in multiple contexts of experience-dependent plasticity.

## AUTHOR CONTRIBUTIONS

E.C.D., M.C.H.B, and M.K.M. conceived and designed the experiments. E.C.D., M.C.H.B, and M.K.M performed all experiments and data analyses in the laboratory of M.K.M. M.K.M. and E.C.D wrote the manuscript. This work was funded by grants to M.K.M. M.K.M initiated and supervised the project.

## ACKNOWLEDGMENTS

This work was supported by the Braude Foundation, NIH MH109341and MH098016 (to M.K.M.), and NS050274 (to Hopkins Neuroscience Multiphoton Facility). We thank C. K. Timmerman for regenerating a key mouse line, M. Pasparakis for GFPRelA and RelA^F/F^ mouse lines, T.R. Gamache for assistance with analysis of p65 amino acid sequence, and J.L. Pomerantz and members of the Meffert laboratory for discussion and critical reading of the manuscript.

